# Wide-field imaging of cortical neuronal activity with red-shifted functional indicators during motor task execution

**DOI:** 10.1101/410365

**Authors:** Elena Montagni, Francesco Resta, Emilia Conti, Alessandro Scaglione, Maria Pasquini, Silvestro Micera, Anna Letizia Allegra Mascaro, Francesco Saverio Pavone

## Abstract

Intracellular concentration of free calcium ions in neuronal populations can be longitudinally evaluated by using fluorescent protein indicators, called genetically encoded calcium indicators (GECIs). GECIs with long emission wavelengths are particularly attractive for deep tissue microscopy *in vivo*, and have the additional advantage of avoiding spectral overlap with commonly used neuronal actuators like Channelrhodopsin.

Here we investigated the performances of selected red-shifted GECIs through an *ex vivo* characterization and *in vivo* imaging of cortical mouse activity during motor task execution. Cortical neurons were infected with adeno-associated virus (AAV) expressing one of the red GECI variants (jRCaMP1a, jRCaMP1b, jRGECO1a, jRGECO1b). First we characterized the transfection in terms of extension and intensity using wide-field fluorescence microscopy. Next, we used RCaMP1a to analyse the cortical neuronal activity during motor behaviour. To that end, wide-field fluorescent microscopy and a robotic device for motor control were combined for simultaneous recording of cortical neuronal-activity, force applied and forelimb position during task execution.

Our results show that jRCaMP1a has sufficient sensitivity to monitor *in vivo* neuronal activity over multiple functional areas, and can be successfully used to perform longitudinal imaging in awake mice.

## 1. Introduction

The calcium ion is widely used in neuronal physiology as a competent indirect reporter of neural activity. One of the main advantages is that concentration fluctuations of this ion in its free form during neuronal activity are among the highest^1-2^.

Genetically encoded calcium indicators (GECIs) have partially replaced both organic sensors and electrophysiological methods since they present considerable advantages, such as the possibility of addressing their expression into specific cellular populations and their stable transfection over time that allows long-term studies of neuronal activity *in vivo*^3-4^.

GCaMP is the most used GECI, since this green emitting sensor presents a fast kinetics of calcium binding, high brightness and great sensitivity^5^. Nevertheless, GCaMP indicator has several limitations due to its excitation and emission spectra. The blue excitation light used in standard fluorescence microscopy can cause photodamage and it is highly scattered in tissue. Furthermore, the green emission of GCaMP is absorbed by hemoglobin^6^, which reduces the penetration depth of imaging in vertebrates *in vivo*. Finally, the GCaMP excitation spectrum overlaps with that of light-sensitive ion channels, such as channelrhodopsin-2 (ChR2), which limits the simultaneous use of green GECIs and optogenetic techniques^7-8^.

These limitations led to an increasing interest in red variant GECIs, namely RGECIs, which are characterized from a structure similar to that of the GCaMP but longer emission wavelengths. RGECIs are composed of a circular permutated thermostable red fluorescent protein (RFP), calmodulin (CaM) and M13, a peptide sequence from myosin light chain kinase^9-10^. Among the most commonly RGECIs used, there are two variants based on two different RFPs: *mRuby* (like in jRCaMP1a and jRCaMP1b) and *mApple* (jRGECO1a and jRECO1b)^8^.

The red-GECIs present multiple advantages for *in vivo* imaging: (i) a longer excitation wavelength which penetrates deeper into tissue due to a reduced scattering by endogenous fluorophores; (ii) red fluorescence is less absorbed than green fluorescence by both endogenous fluorophores and hemoglobin in mammalian tissue^8-11^.

In the last years, different fluorescent sensors and optogenetic tools have been combined to achieve simultaneous optical manipulation and recording of neuronal activity, but each association had some limitations in use. For example, the coupling of red-shifted organic voltage-sensitive dye (VSD) as sensor, and ChR2 as activity manipulator, is limited by fast degradation and photoswitching of VSD^12^. At the same time, the combination of GCaMP6s and red-shifted C1V1 was limited by C1V1 opsin photoswitching when illuminated by blue light^13^. On the other hand, R-GECIs are more easily compatible with ChR2 activation for the purpose of simultaneous neural circuits activity monitoring and manipulation over time^14^.

Currently, the most flexible approach for inducing the expression of GECI is viral transfection, that allows targeting a specific cellular population in selected functional areas. Then, the fluorescence emission of GECI can be recorded *in vivo* using non-invasive imaging techniques like wide-field fluorescence microscopy (WFFM). This technique allows the registration of neuronal activity on distant functional cortical areas over both cortical hemispheres with millisecond resolution^15^.

In order to choose the best red-shifted GECI for long-term *in vivo* studies using one-photon microscopy, here we examined four different commercially available RGECIs in terms of both intensity and extension of the transfection. Finally, we show the application of jRCaMP1a to the monitoring of neuronal activity during the execution of a motor task.

## 2. Methods

### Animals

All mouse handling and manipulations were performed in accordance with the rules of the Italian Minister of Health. In this study, we used both male and female C57BL mice (age > 1 years). All the animals included in this study were housed in an animal room with a 12-hour/12-hour light/dark cycle, with food and water available and unlimited.

### Viral injection and optical windows

All surgeries were conducted under isoflurane anesthesia (1.5–2%) and local anesthetic lidocaine 2% was administrated as necessary.

For the viral injection, the animals were placed into a stereotaxic apparatus and both the skin over the skull and periosteum were removed. After that, a small hole was thinned into the skull (Ø 0,4mm) on the right hemisphere using a dental drill.

For every indicator (n_mice_ = 4), we injected 250nl at two different depths 0.4mm and 0.8mm from the dura, into the somatomotor cortex (+1.5mm mediolateral and −1.5 anteroposterior from bregma).

The injection was made through use of a capillary (Ø of the tip: 50µm) connected to Picospritzer (Picospritzer III – Science Products™).

The virus used were:

### jRCaMP1a

pGP-AAV-syn-NES-jRCaMP1a-WPRE.211.1488

### jRCaMP1b

pGP-AAV-syn-NES-jRCaMP1b-WPRE.211.1519

### jRGECO1a

pGP-AAV-syn-NES-jRGECO1a-WPRE.111.1670

### jRGECO1b

pGP-AAV-syn-NES-jRGECO1b-WPRE.111.1721

The AAV9 serotype (AAV9) allowed indicator expression at central nervous system level. Moreover, the neuro-specific promoter (synapsin) targeted both excitatory and inhibitory neurons^16^ while NES motif limited expression to the cytoplasm.

In order to study the cortical activity during motor task execution, we injected the primary motor cortex (+1.75mm mediolateral and +0.5mm anteroposterior from bregma) of 3 mice with the jRCaMP1a construct (500nl) at 0.5mm cortical depth.

In addition, all mice were implanted with a cover glass to allow free optical access to the cortex. A custom-made aluminum head-bar was attached to the skull to allow the fixation of the head during imaging analysis. Dental cement (Super Bond C&B – Sun Medical) was used as fixative for both surgeries.

### Wide-field fluorescence microscope

The custom-made wide-field imaging setup was equipped with an excitation source for imaging of RGECIs fluorescence (595nm LED light, M595L3 Thorlabs, New Jersey, United States) and a pass-band filter (578/21nm, Semrock, Rochester, New York USA) allows the selection of the excitation wavelength. A dichroic filter (606nm, Semrock, Rochester, New York USA) above the objective (2.5x EC Plan Neofluar, NA 0.085) deflected the light beam. A 3D motorized platform (M-229 for xy plane, M-126 for z-axis movement; Physik Instrumente, Karlsruhe, Germany) allowed animal positioning under the objective.

The fluorescence signal was selected by a pass-band filter (630/69, Semrock, Rochester, New York USA) and focused by a focal lens (500mm) on the sensor of a high-speed complementary metal-oxide semiconductor (CMOS) camera (Orca Flash 4.0 Hamamatsu Photonics, NJ, USA) where 512 by 512 px^2^ images covering 4,4 by 4,4 mm^2^ of cortex were acquired.

### Wide-field calcium imaging

Calcium imaging sessions were performed from 2 to 4 weeks after surgery in a resting state condition (the animals were awake, but they were not subjected to stimuli). The imaging field was manually located using reference images of previous recording days at the beginning of each session.

During the last week, one experimental group (jRCaMP1a, injection in motor cortex, n=3) was subjected to motor training for 5 consecutive days.

#### Data analysis

The analysis have been performed using ImageJ, OriginPro and Mesoscale Brain Explorer (MBE) software. Each experimental session consisted in 40 second of recording (exposure time: 40ms). The fluorescence traces were analysed both on the whole acquired field and on five regions of interest (ROIs) of 30 pixel in size, representative of specific functional areas: primary and secondary motor cortex (M1 and M2), primary sensory cortex in barrel field and forelimb region (S1BL and S1FL) and retrosplenial cortex (RS) (*fig 4B*).

For each week, we calculated the average ΔF/F value on the whole acquired field, according to the following formula:

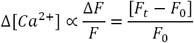

Where ***F***_*0*_ was the average of baseline fluorescence intensity and ***F***_*t*_ was the fluorescence issued at a given time. We only considered fluorescence peaks with intensity 20% higher than the baseline. The threshold value was calculated for every week as the average of the intensity in a ROI (0,24 mm^2^) far from fluorescence region.

#### Full width half-maximum

For each indicator, we chose three frames corresponding to the maximum fluorescence peak in the ***time*** at the fourth week (exposure time 40ms, LED power 23mW). On these frames we calculated the full width at half maximum (FHWM) of fluorescence profiles, tracing two straight lines passing through the injection site (rostro-caudal and medium-lateral plan). Moreover, on the same frames, we obtained the max peak amplitude of fluorescence profiles as average ΔF/F values (*fig 2*).

**Figure 1:**
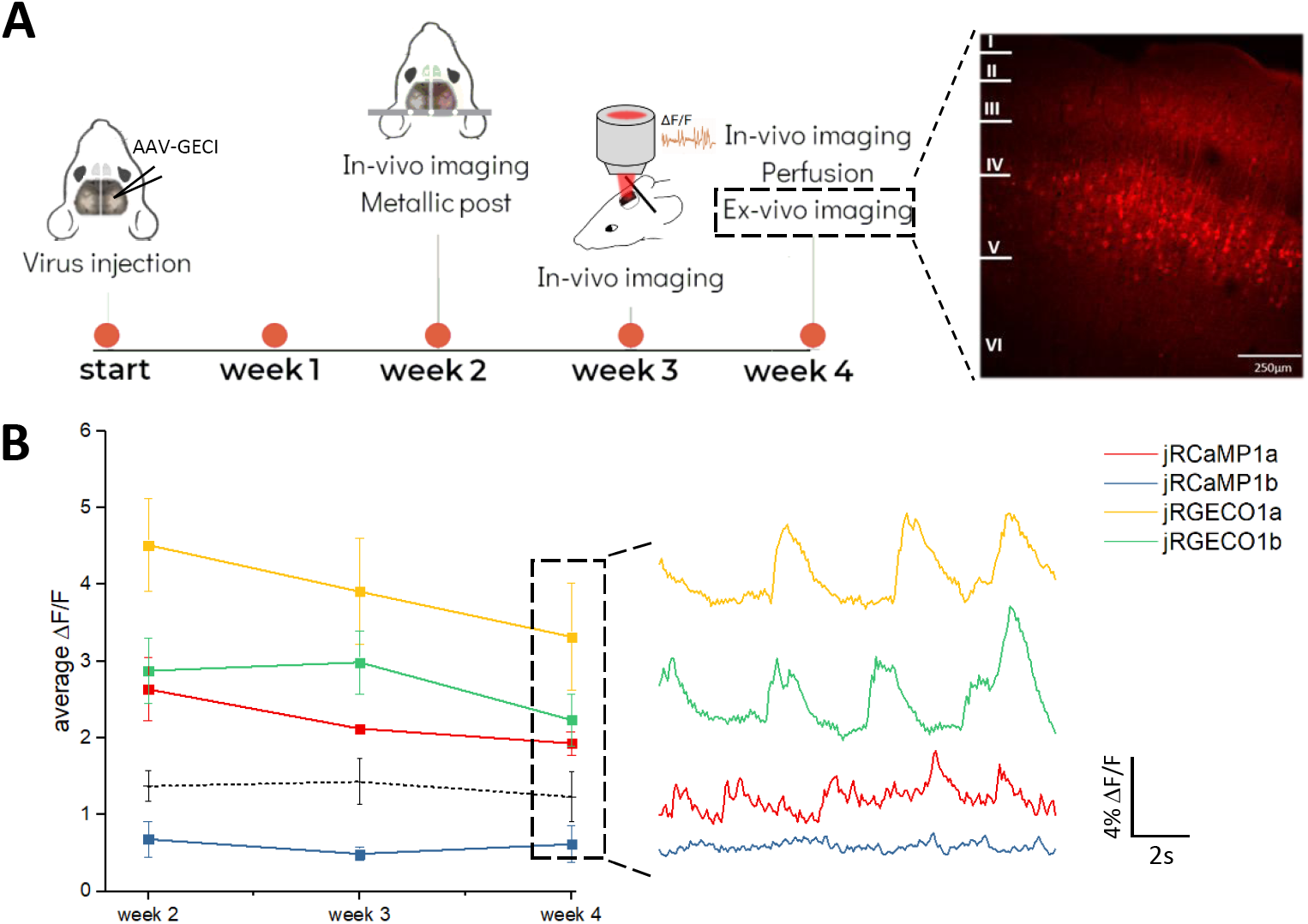
Stability of red-shifted indicators expression over 4 weeks. **(A)** Experimental timeline: first, the injection of AAV-GECI is performed at cortical level. After 2 weeks, a metallic post was implanted and the in **<u>vivo</u>** imaging sessions started, which were repeated weekly. After 4 weeks, brain slices were analysed by wide-field microscopy. **(B)** Average cortical activity in vivo along 4 weeks (n_mice_= 4 exp-group. **jRCaMP1a**: ΔF/F_W2_= 2,6±0,4, ΔF/F_W3_= 2,1±0,1 ΔF/F_W4_= 1,9±0,1. **jRCaMP1b**: ΔF/F_W2_= 0,6±0,2, ΔF/F_W3_= 0,5±0,1, ΔF/F_W4_= 0,6±0,2. j**RGECO1a:** ΔF/F_W2_= 4,5±0,6, ΔF/F_W3_= 4±0,1 ΔF/F_W4_= 3,3±0,7. **jRGECO1b:** ΔF/F_W2_= 2,9±0,4, ΔF/F_W3_= 3±0,4 ΔF/F_W4_= 2,2±0,3. **Threshold:** ΔF/F_W2_= 1,3±0,2, ΔF/F_W3_= 1,4±0,3 ΔF/F_W4_= 1,2±0,3). On the right, example traces of in-vivo recorded fluorescence activity for every indicator (during the fourth week). Values are reported as average±SEM.

**Figure 2:**
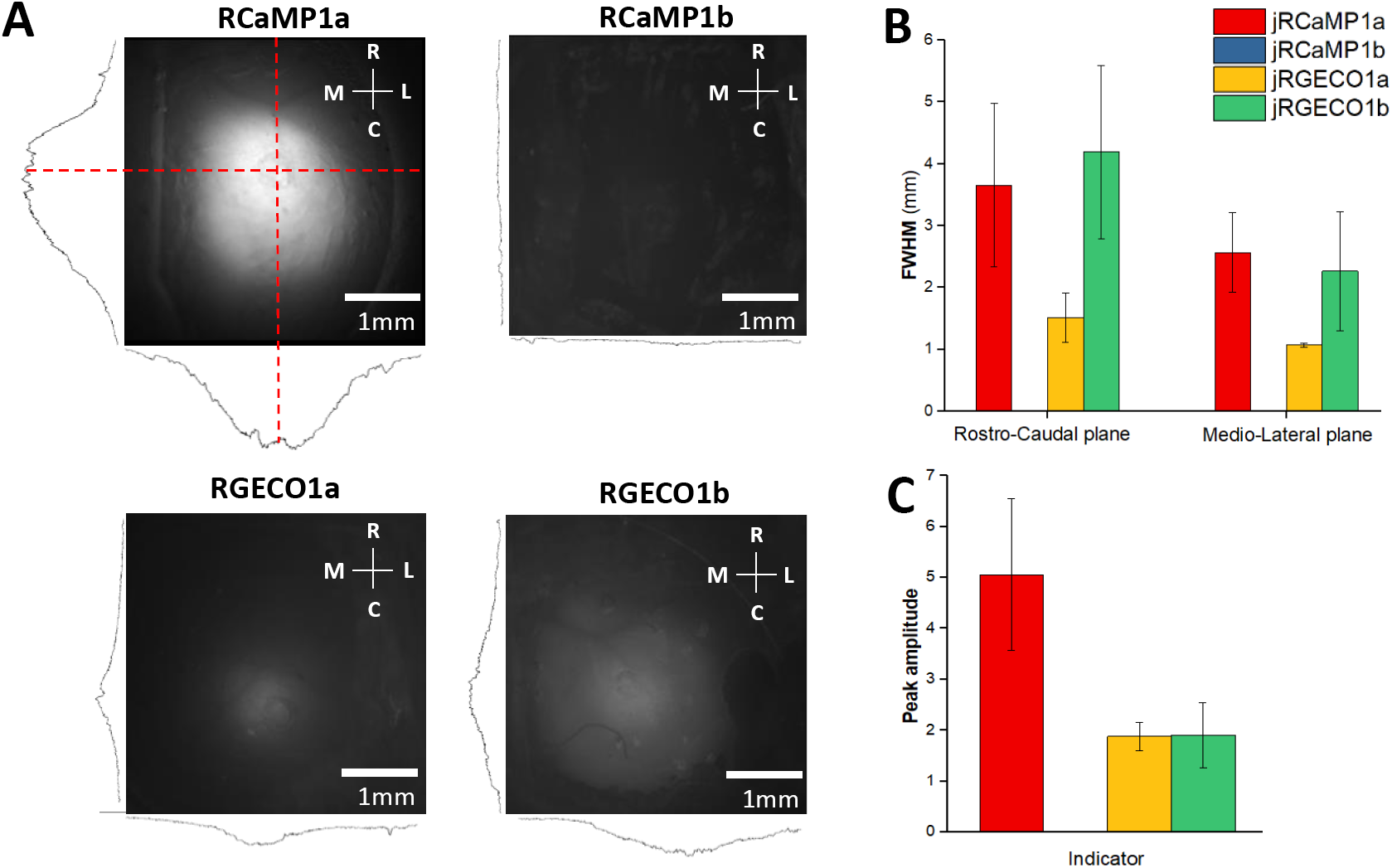
In-vivo quantification of AAV transfection of red-shifted calcium indicators. **(A)** Representative images in-vivo of spatial distribution of each indicator. On the bottom and on the left of each image, the fluorescence profiles along both the medium-lateral and rostro-caudal plane are reported. **(B)** Full width at half maximum (FWHM) of fluorescence profiles for each indicator (n_mice_= 4 exp-group) along both rostro-caudal (left, FWHM-RC_jRCaMP1a_= 3,7±1,3mm, FWHM-RC_jRGECO1a_= 1,5±0,4mm, FWHM-RC_jRGECO1b_= 4,2±1,4mm) and medio-lateral plane (right, FWHM-ML_jRCaMP1a_= 2,6±0,6mm, FWHM- ML_jRGECO1a_= 1,1±0,1mm, FWHM-ML_jRGECO1b_= 2,3±1,0mm). jRCaMP1b fluorescence is comparable to noise values. **(C)** Average ΔF/F values of fluorescence peaks at the transfection site 4 weeks after AAV injection (ΔF/F_jRCaMP1a_= 5.1±1.5, ΔF/F_jRGECO1a_= 1.9±0.3, ΔF/F_jRGECO1b_= 1.9±0.5). Values reported as average ± SEM. Scale bar, 1mm.

#### *Ex vivo* imaging

Four weeks after injection, mice were perfused with 20-30ml of 0.01M PBS (pH 7.6) and 150ml of 4% paraformaldehyde (PFA). After the perfusion, we obtained brain coronal slices (100µm thick) by use of vibratome (Vibratome Series 1500 – Tissue Sectioning System). On each slice, we have studied the rostro-caudal transfection extension using wide-field fluorescence microscopy (*fig 3A*, exposure time 12ms, LED power 23mW).

**Figure 3:**
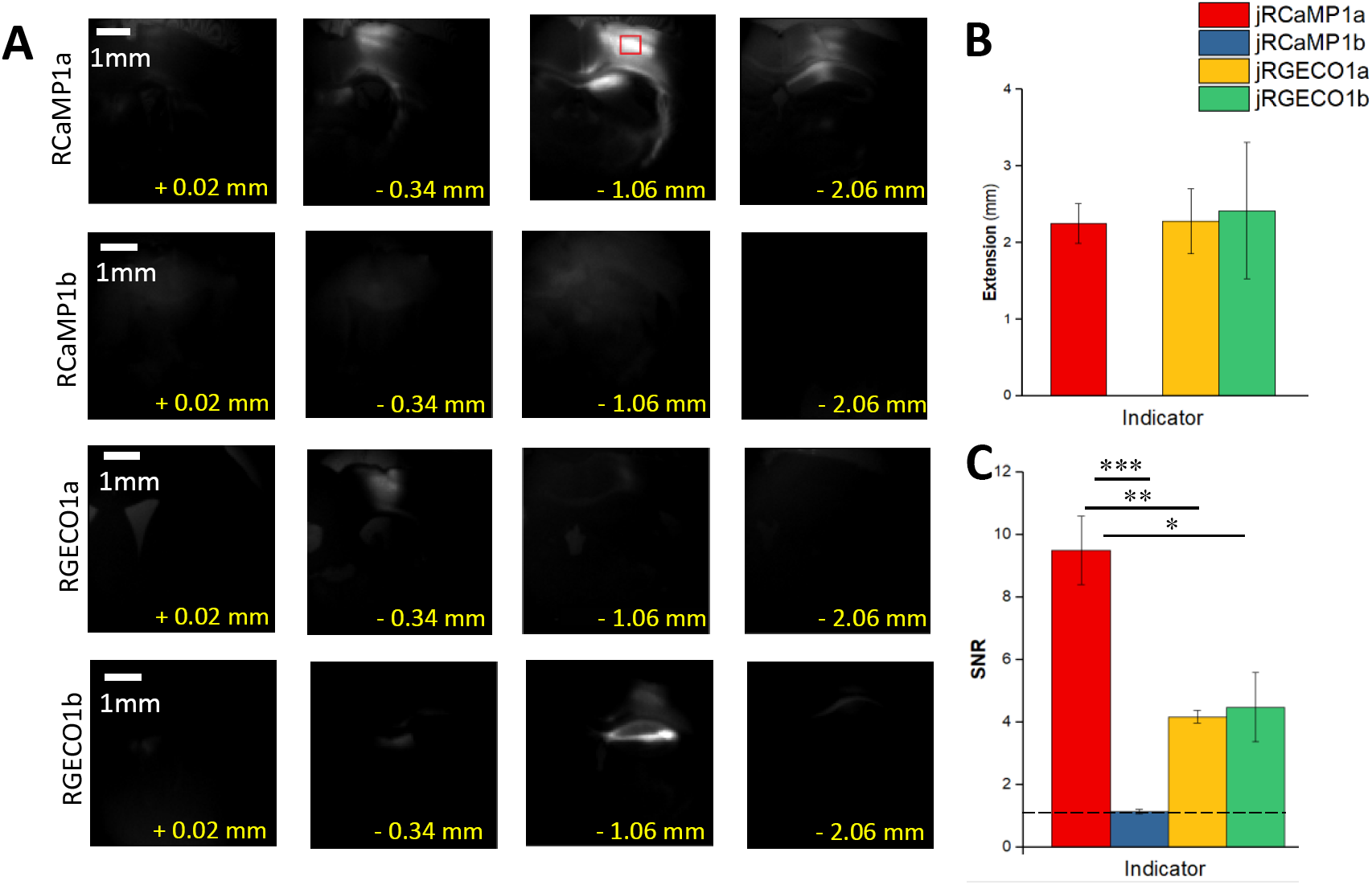
Ex-vivo characterization of red shifted calcium indicators. **(A)** Representative image sequences of rostro-caudal extension of brain slices showing rostro-caudal extension of transfection for each indicator. The images are selected at the same distance from bregma (from +0.02mm to −2.06mm relative to bregma) one month after AAV injection‥ **(B)** Rrostro-caudal extension of transfection (n_mice_=4 exp-group), EXT_jRCaMP1a_= 2,6±0,2mm, EXT_jRCaMP1b_= 0mm, EXT_jRGECO1a_= 2,3±0,4mm, EXT_jRGECO1b_= 2,8±1,2mm). **(C)** Signal-to-noise ratio for each indicator (SNR_jRCaMP1a_= 9,4±1,1, SNR_jRCaMP1b_= 1,1±0,1, SNR_jRGECO1a_=4,2±0,2, SNR_jRGECO1b_= 4,5±1,1). One-way ANOVA followed by the Bonferroni test: ***P_(RCaMP1a/RCaMP1b)_=3*10^−5^; **P_(RCaMP1a/RGECO1a)_=0.001; *P_(RCaMP1a/RGECO1b)_=0.003. Values reported as average±SEM. Scale bar, 1mm.

In addition, on the three brightest slices, we have evaluated signal to noise ratio (SNR, *fig 3C*) calculated as follows:

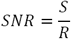

Where **S** was the fluorescence intensity averaged on a 0,24mm^2^ ROI centered on the brightest area, mediated on the three brightest slices, while **R** was the mean intensity of noise obtained in the same slices but in an area furthest from the transfection site (∼ 2 mm).

### Robotic platform

Mice expressing jRCaMP1a in motor cortex performed 5 consecutive days of training in the robotic device (Pasquini *et al.*, under review). The single daily session consisted in 15 cycles or “trials” of active retraction of left forelimb associated with a sweetened condensed milk reward (10µl) at the end of each task. Each day, before the training, we recorded the “baseline”, which was the spontaneous neuronal activity in resting state (40s of acquisitions, 40ms of exposure time).

The robotic device (M-platform) is based on the one described in *Spalletti et al.*^17^. It is composed of a linear actuator, a 6-axis load cell (Nano 17, ATI Industrial Automation, USA), a precision linear slide with an adjustable friction system and a custom-designed handle where the left wrist of the mouse was allocated, which allowed a transfer of the force applied by animals to the sensor.

In each trial, first a linear motor pushed the slide and extended the mouse left forelimb by 10mm (passive phase). Then, the motor was quickly decoupled form the slide and the mouse was free to voluntarily pull the handle back (active phase). Two acoustic signals informed the mouse of the end of passive phase (0.5s) and the reaching of a target position (1s), which was associated with reward.

During the exercise, the robotic device integrated in the wide-field microscope allows simultaneous recording of three different data associated with each other (fig 4A): (i) the recording of cortical activity as a change in fluorescence signal, (ii) the force applied by left forelimb situated in a handle and (iii) the limb position, estimated by the movement of the slide using an IR position sensor located on the slide and recorded by an IR camera (EXIS WEBCAM #17003, Trust).

**Figure 4:**
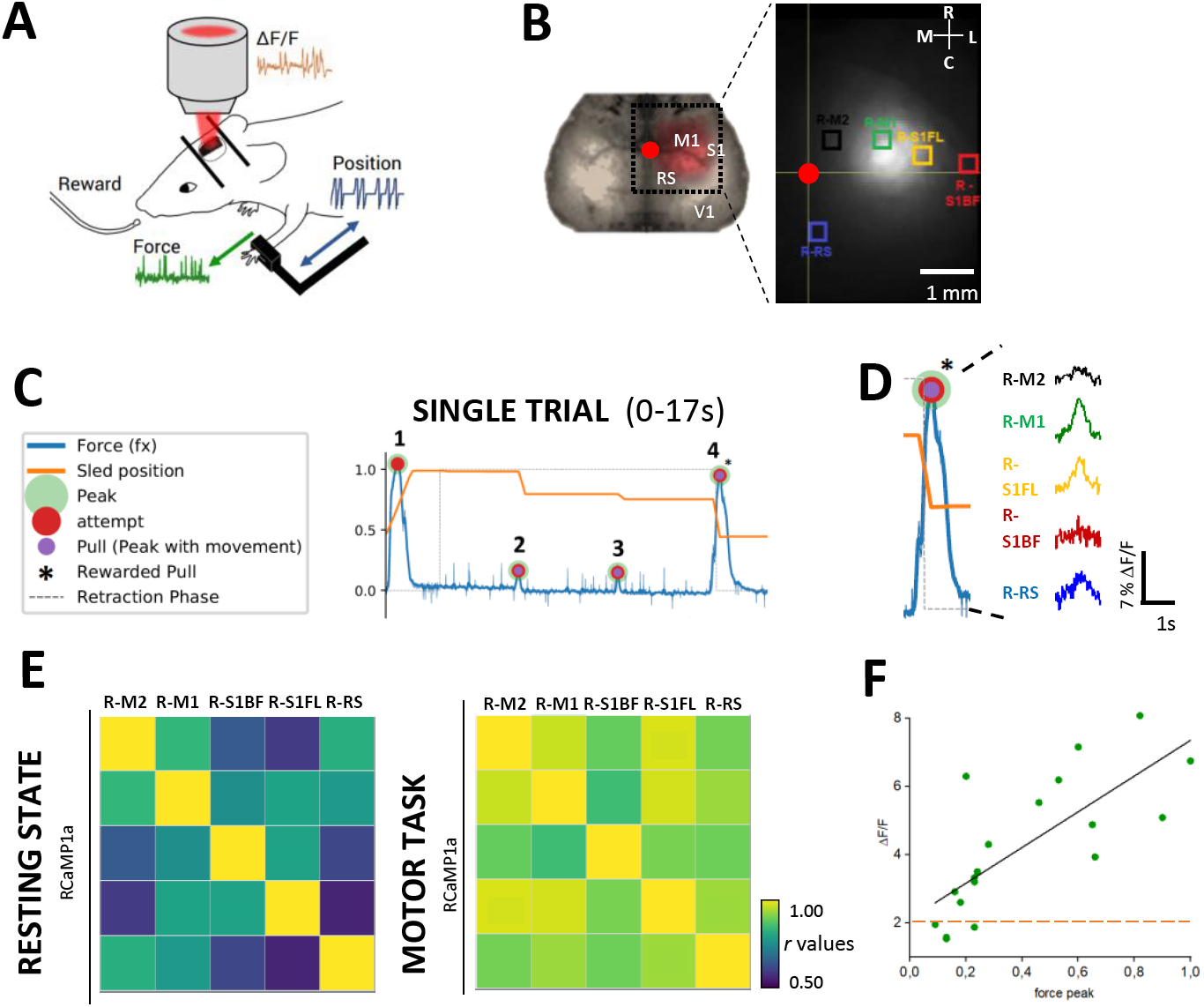
Cortical neuronal activity in resting state and during execution of motor task. **(A)** Schematic view of the M-platform used for training integrated with the wide-field microscope, which allows simultaneous recording of three different data: the force applied by forelimb (green), the position of left forelimb (blue) and the contralateral cortical activity as a change in the indicator fluorescence intensity (red). **(B)** Schematic of field of view of the microscope, highlighted by the black dotted square, superimposed to the cortical hemispheres. The 5 cortical functional areas used for correlation matrix are reported (inset). The ROIs are primary and secondary motor cortex (M1 and M2, green and black respectively), primary sensory cortex in both barrel field and forelimb region (S1BF and S1FL, red and yellow respectively) and retrosplenial cortex (RS, blue). Bregma in reported as red spot. **(C)** Example of force trace obtained by the load cell on the M-platform during a single trial in the forelimb active retraction phase. The forelimb position is in orange, and the force applied by mouse in blue. The force peaks trace shows the attempts to move the slide (red spot), the force peaks associated with movement of the slide (purple spot) and the pulling associated with reward (black star). **(D)** Example of neuronal activity recorded simultaneously on the 5 cortical areas during a single force peak (blue) which is associated with the movement of forelimb (orange) and the rewarded pull (black star). **(E)** Linear average correlation matrices between five selected ROIs, obtained both during resting state condition (left, n_trials_= 12) and motor task execution (right, n_trials_=12) in the same mouse. **(F)** Scatter distribution of the relationship between the maximum fluorescence activity of M1 and the corresponding force peak (n_mice_=1, n_peak_=19). The black line shows the best fit (intercept= 2,1±0,5, slope=5±1). Orange dashed line reports the threshold measured as the average of M1 maximum activity in resting state condition in the same mouse (n_peak_=19; ΔF/F=2,06). Values reported as average±SEM.

#### Scatter distribution

We selected the M1 maximum activity associated with force peaks exerted (n_peak_=19). The threshold was obtained averaging the intensity of maximum peaks during resting state condition in the same mouse (n_peak_=19) (*fig. 4F*).

### Correlation Matrix

These correlation matrixes were realized using data from a single *mouse* and analyzed by mesoscale brain explorer (MBE) program^18^.

The correlation matrixes were produced calculating the correlation index (r) of cortical activity (*Δ*F/F) between the five *functional* areas.

*Primary* motor cortex (M1) is a functional brain area highly involved in motor control^19^.

Somatosensory (S1) and M1 cortices are reciprocally connected, and so sensory feedbacks have been reported to play an important role in driving learned movements^20-21^. This condition led the selection of S1 in forelimb region (S1FL).

Secondary motor cortex (M2) receives several sensory afferences and has reciprocal connection with retrosplenial cortex. It contributes to functional recovery after stroke in primary motor cortex and it has a predictive role to drive the motor behavior. Moreover, M2 plays a role into the neuronal circuit mechanism of voluntary actions ^22-23-24^.

Retrosplenial cortex (RS) supports spatial working memory, but its projection to M2 suggest a role in motor control and sensorimotor integration too^22-25^.

Conversely, sensory cortex in barrel-field region (S1BF) was chosen as a counter-test because it is principally involved in whisker-dependent behaviours and should not be primarily involved in lever pulling task^26^.

The correlation index depicts how the activity in one area correlates with other areas in the matrix, and was influenced by both (i) fluorescence intensity variation and (ii) time delay with which the same peak occurs in two different areas.

The correlation matrix (fig 4E) was calculated on 12 of 15 consecutive motor trial. We selected imaging datasets were satisfying a specific condition, which was when the peak force necessary to pull the slide in an attempt managed to produce displacement of the slide (peak 2 and 3 in fig 4C) and when it was associated with a reward (rewarded pull, peak 4 in fig 4C). We discarded the trials where the paw slipped away from the slide.

On the same mouse, the correlation matrix during resting state condition was obtained on 12 different baseline imaging datasets.

## 3. Results

### Experimental timeline

We characterized four different red-GECIs (jRCaMP1a, jRCaMP1b, jRGECO1a, jRGECO1b) locally expressed at somatomotor cortex level by using *in vivo* and *ex vivo* fluorescence imaging.

To this aim, we performed intraparenchimal injections of adeno-associated viruses expressing one of the red GECI variants under the human synapsin1 (neuron-specific) promoter (AAV-SYN1-red GECI variant). In order to evaluate both the stability of transfection over the weeks and *in vivo* indicator distribution on the cortex, an imaging session was performed at 2, 3 and 4 weeks after surgery. To quantify more precisely the extension of the transfection, we performed *ex vivo* imaging on brain slices four weeks after the injection (*fig 1A*).

### *In vivo* wide-field calcium imaging over 4 weeks

During the imaging session *in vivo* we recorded the spontaneous cortical activity to evaluate the sensitivity of the indicators in a resting state condition (*fig 1B)*.

Next, we performed a comparison of the average cortical activity between each week. Our 29results showed that only jRCaMP1b indicator had a very low expression level, which was comparable to the background noise. On the other hand, among the other three indicators, jRGECO1a had the best average values (*fig 1B*).

We showed that the indicator expression was stable over time (fig 1B). This condition allows long-term studies of neuronal activity.

### *In vivo* fluorescence distribution in the space of four R-GECIs

Four weeks after injection, on the brightest frames, we quantified *in vivo* fluorescence extension in the space on both rostro-caudal and medium-lateral plane (*fig 2A*). To this aim we evaluated the full width at half maximum (FWHM) and the peak amplitude of fluorescence profile for each indicator.

The peak amplitude (*fig 2C*) gives an indirect estimation of indicator sensitivity and concentration *in situ*. Our results showed that jRCaMP1a was the brightest near the injection site.

Moreover, we investigated the FWHM (*fig 2B*), which is an index of indicator distribution in space. jRCaMP1a and jRGECO1b exhibited the largest *in vivo* fluorescence extension along both planes. Since jRGECO1b did not exhibit high expression level (*fig 2C*), we identified the jRCaMP1a indicator as the best in terms of *in vivo* distribution in space, sensitivity and concentration level.

Furthermore, we found that indicator expression was isotropic from the injection site along both planes for all the indicators (*fig 2B*).

### *Ex vivo* characterization of R-GECIs transfection

To finely quantify the expression profile of the indicators throughout the cortex, we further assessed both the rostro-caudal extension and the expression level of transfection *ex vivo*.

Our results confirmed that three out of four indicators were successfully transfected (fig 3A), while the expression level of jRCaMP1b indicator was not detectable in our analysis.

Compared to the other indicators, jRCaMP1a construct showed both a widest transfection at cortical level (fig 3B) and the best signal to noise ratio (fig 3C), which can be associated with an higher in-situ expression level.

Taken together, these results allowed us to identify the best red-shifted calcium indicator for wide-field imaging: jRCaMP1a, which is characterized by both wide *in vivo* fluorescence distribution and high sensitivity.

### Evaluation of jRCaMP1a sensitivity to neuronal activity during motor task execution

We further investigated the performance of jRCaMP1a on awake mice during motor behavior. We transfected jRCaMP1a indicator at motor cortex level and monitored neuronal activity 4 weeks after the injection on five different functional areas during motor task execution on a robotic device (M-platform^19^).

The robotic platform allowed us to record simultaneously the force applied by forelimb, the position of left forelimb and the contralateral cortical activity (*fig 4A*).

The motor task consisted of 15 consecutive active retractions of left forelimb; after reaching the target (fully-retracted) position, the mouse received a milk reward. This training session was performed for 5 days.

The correlation matrices were obtained during both resting state condition and motor task execution (*fig 4E*). In the last case, we selected only the timeframes of the imaging dataset corresponding to force peaks associated with forelimb movement (peak 4 in *fig.4C* and *fig. 4D*).

Our results showed a greater and more widespread correlation of neuronal activity during the motor task compared to resting state. The higher variation of correlation index takes place between two ROIs pair: R-M2/R-S1FL and R-RS/R-S1FL (*fig 4E*).

Moreover, we analyzed the potential of our paradigm in detecting the neuronal activation in motor cortex associated with a broad range of forces applied. We found that when the force applied during the active retraction by the forelimb has small intensity, the M1 activity associated is detectable and slightly higher than the average value measured in resting state condition. As the applied force increases, the maximum activity of M1 increases linearly (*fig 4F*).

These results allowed us to conclude that the extension of the transfection was sufficient to simultaneously record the activation of several cortical areas distant from injection site, and that the jRCaMP1a is sufficiently sensitive to report the variation of neuronal activity associated with the modulation of applied forces.

## 4. Discussion

In this study, we performed *in vivo* and *ex vivo* characterizations of the expression patterns of four different red-shifted GECIs transfected at cortical level. We confirmed that indicator expression is stable over time for all sensor except for the jRCaMP1b indicator, which showed transfection level comparable to noise in each analysis performed. On the other indicators, we show that there is a slight (statistically non-significant) reduction of average ΔF/F over the weeks. Although the mice were acclimated to the environment for few minutes before each imaging session, this decrease could be attributed to changes in the emotional state of the animals over the days and the weeks.

jRGECO1a indicator (mApple based) showed the highest response amplitude over the weeks, in agreement with previous studies showing that it is the most sensitive indicator with fastest rise kinetics^8^. Nevertheless, it has been shown that both mApple based-indicators (jRGECO1a and jRGECO1b) are affected photoswitiching effect when illuminated by blue light^8-27^, thus limiting their use for optogenetic studies where blue activated opsin are used.

The *in vivo* and *ex vivo* quantification of indicator distribution in the cortical space showed jRCaMP1a as the best sensor in terms of sensitivity and signal to noise ratio. Our results are in agreement with previous two-photon studies where jRCaMP1a is identified as the brightest indicator in the calcium-bond state^8^. In addition, we found that jRCaMP1a construct is associated with the widest rostro-caudal extension of transfection four weeks after injection. jRCaMP1a expression involved several well-defined functional areas along the right hemisphere, allowing in combination with wide-field microscopy the simultaneous study of the functionality of five cortical areas.

In conclusion, although jRGECO1a has been previously identified as the most sensitive and with faster rise kinetics compared to the other indicators^8^, our results indicated jRCaMP1a construct as the more suitable for *in vivo* wide-field imaging studies.

We therefore choose jRCaMP1a indicator for the subsequent study of neuronal activity during a motor exercise performed on a robotic device.

We demonstrated that our approach allowed investigating the interplay between the selected functional areas during an active behavior involving movement of the limb. In agreement with previous studies, we showed the involvement of sensorimotor-areas in movement control. The barrel-field region showed (i) the lowest correlation degree with the other functional areas during the exercise and (ii) the lowest variation of correlation index with R-M1 between resting state and motor task execution. On the other hand, we showed an extensive increase in correlation of neuronal activity during the task compared to resting state for each ROIs pair.

In conclusion, by non-invasive wide-field imaging studies we demonstrated that jRCaMP1a transfected in one cortical hemisphere allows simultaneous recordings of the activity in several functional areas. Furthermore, this combination of tools is capable of detecting the variation of neuronal activity at motor cortex level associated with a broad ranges of applied forces.

Finally, we anticipate that the well-defined excitation and emission spectra of ChR2 and jRCaMP1a will enable simultaneous all-optical manipulation and wide-field recording of neuronal activity in awake animals. Our approach will be targeted to the investigation of optogenetically induced neuronal activation associated with complex movements. The clear calcium response evoked by ChR2 stimulation will allow to better understand neural processes underlying specific behaviors.

## Acknowledgements

This project has received funding from the European Union’s Horizon 2020 Research and Innovation Programme under Grant Agreement No. **720270** (HBP SGA1). In addition, it was supported by the European Union program H2020 EXCELLENT SCIENCE – European Research Council (ERC) under grant agreement ID n.**692943** (BrainBIT) and by Regione Toscana PAR-FAS 2007-2013 Bando Salute 2014 - RONDA (Robotica indossabile personalizzata per la riabilitazione motoria Dell’arto superiore in pazienti neurologici).

